# Helminth ecological requirements shape the impact of climate change on the hazard of infection

**DOI:** 10.1101/2023.09.11.557173

**Authors:** Chiara Vanalli, Lorenzo Mari, Renato Casagrandi, Marino Gatto, Isabella M. Cattadori

**Affiliations:** Center for Infectious Disease Dynamics and Department of Biology, The Pennsylvania State University, University Park, 16802 PA, USA; Dipartimento di Elettronica, Informazione e Bioingegneria, Politecnico di Milano, 20133 Milano, Italy

**Author notes:** Corresponding author: Chiara Vanalli (. **Authorship statement** CV performed the literature analysis, modeling work, result visualization and wrote the first draft of the manuscript. LM, RC and MG guided the modeling development. IMC developed the original idea, advised on the analysis, results interpretation and contributed to the first draft writing. All authors contributed substantially to manuscript drafting and editing.

**Keywords:** Trichostrongylidae, Free-living stages, Hatching, Development and mortality, Seasonality, Spatial distribution, Co-circulation

## Abstract

Outbreaks and spread of infectious diseases are often associated with seasonality and changes caused by global warming. Free-living stages of soil-transmitted helminths are highly susceptible to environmental drivers, however, how multiple climatic variables affect helminth species, and the long-term consequences of these interactions, is poorly understood. We used experiments on nine trichostrongylid species to develop a temperature- and humidity-dependent model of infection hazard, which was then implemented at the European scale under climate change scenarios. Intestinal and stomach helminths exhibited contrasting climatic responses, with the former group strongly affected by temperature while the latter primarily impacted by humidity. These differences generated seasonal changes in the timing and intensity of the infection hazard and spatial heterogeneities within and between the two groups. A future range expansion of both groups toward northern latitudes is expected to create new opportunities for the co-circulation of the studied helminth species.

## Introduction

The many forms of disruption associated with climate change, like warming temperature, extreme climatic events or shifts in climatic ranges, are expected to strongly affect ectotherm species by selectively targeting components of their life cycle and dynamics (Deutsch et al., 2008; Paaijmans et al., 2013; Wagner et al., 2023). These climatic changes are also predicted to alter the severity and spread of many circulating infections, whose transmission depends on the survival and development of stages that live free in the environment, as is the case of soil-transmitted helminths with direct life cycle (Hoar et al., 2012; Molnár et al., 2013b, 2017; Rose et al., 2016; Smith, 1990), as well as of those parasites that require intermediate invertebrate hosts or vectors for maturation and transmission (Carraro et al., 2017; Kutz et al., 2002, 2005a; Molnár et al., 2013a; Schjetlein and Skorping, 1995). Assessing the net effect of climate warming on parasite transmission has proved to be challenging because of the non-linear thermal responses that parasite traits often exhibit (Gehman et al., 2018; Molnár et al., 2017). Indeed, the trade-off of faster development but lower survival commonly observed for parasites (Hoar et al., 2012; Rose et al., 2016; Smith, 1990) and vectors (Mordecai et al., 2019, 2017) exposed to increasing temperatures suggests that transmission is far from being linearly related to temperature, but more likely the result of complex interactions with asymmetric humped-shaped thermal responses. Therefore, the assumption that warming will aggravate the prevalence and severity of current endemic infections cannot be fully generalized among parasite species, and needs to be carefully evaluated across a broad range of temporal and spatial settings (Lafferty and Mordecai, 2016).

Seasonality is one of the strongest environmental forces that affects the fluctuation of parasite prevalence and abundance over time (Altizer et al., 2006; Dowell, 2001; Harvell et al., 2002). Temperature warming could increase or decrease the intensity of parasite transmission without affecting the seasonal trend, alternatively, it could impact both the magnitude and duration of transmission and thus its seasonal shape (Altizer et al., 2006). This latter scenario has been proposed for a few helminth species under climate change, whose current season of spring-to-fall transmission is expected to split into two separate periods, spring and fall, divided by the emergence of a summer minimum (Altizer et al., 2013). For example, a bimodal pattern has been described for the sheep helminth *Haemonchus contortus* in southern Europe (Rose et al., 2016) and for the reindeer nematode *Ostertagia gruehneri* in the artic Canada (Hoar et al., 2012; Kutz et al., 2014; Molnár et al., 2013b) where temperatures have been found to exceed the parasite thermal optimum. In addition to seasonal changes, we should also expect the geographical expansion of parasites in those areas where climate will become more suitable for survival and persistence, but a range contraction where parasites have limited thermal tolerance or slow adaptation to rapid climate changes (Kafle et al., 2020; Kenyon et al., 2009; Kutz et al., 2013, 2009; Short et al., 2017). Importantly, given the strong heterogeneity of global warming expected across geographical areas in the coming decades, these spatial trends should be more apparent at the extremes of the parasite range of distribution, where life history traits are under stronger constraints.

While studies on parasites and vectors suggest that climate warming can alter the seasonal profile of infectious disease transmission and the range of species distribution, when and where changes in these patterns arise, what climatic variables affect these trends, and whether the observed patterns will be maintained as climate becomes warmer needs careful investigation. The frequent assumption is that many of the relationships between parasite life-history strategies and climate can be generalizable across species. However, parasite species might have different thermal responses to the same demographic trait and it is important to examine how these responses change among species or groups of species that have distinct ecological requirements (Molnár et al., 2013a). Moreover, given that natural and domestic animal populations are often infected by a community of parasite species, evaluating whether climate change will promote or prevent opportunities for co-circulation of multiple species and risk of co-infection remains an overlooked issue (Clerc et al., 2018; Graham et al., 2007; Johnson and Buller, 2011). In fact, although these patterns ultimately depend on the distribution and abundance of the host populations, climate modulates the dynamics of free-living stages, including the viability of infective stages and thus the hazard of infection over time and space (Dobson et al., 2015; Molnár et al., 2017; Rose et al., 2016).

In this study, we focused on the most common soil-transmitted helminths of some of the most familiar domestic and wild herbivore species and examined the historical and future impact of climate on the hazard of infection and related life history traits (i.e. egg hatching and mortality, larval development and survival) of stages free-living in the environment. Our goal is to identify commonalities and dissimilarities in the temporal and spatial demography of helminth species with diverse life histories under the direct effect of temperature and relative humidity. Given our interest in the free-living stages, and to reduce the complexity of using several host species, we did not explicitly address the role of the host populations or additional anthropogenic factors, like anthelminthic treatment or livestock management, on the hazard of infection. We focused on disentangling the climatic dependence of rates that drive the dynamics of free-living stages, not of their absolute abundance, making our framework easy to compare with other host-helminth systems. We selected parasites from the Trichostrongylidae family, which includes the genera *Trichostrongylus, Haemonchus, Ostertagia, Cooperia*, and *Nematodirus*, and commonly infect a large number of herbivore species, such as sheep, goats, and cattle for livestock as well as rabbits and hares for wildlife (Anderson, 2000). These helminths have a direct life cycle where infection occurs by ingestion of infective stages available on the pasture; adults colonize the gastrointestinal tract of their hosts, either the intestine or the stomach, and shed eggs in the environment via host’s feces. These helminths represent classical examples of climate dependency identified through laboratory studies (Dobson et al., 2015; Hoar et al., 2012; Kafle et al., 2020; Kutz et al., 2005b; Molnár et al., 2017, 2013b; Rose et al., 2016). In addition, the Trichostrongylidae family is among the ones that cause consistent economic costs to the livestock industry (Charlier et al., 2020), and a better understanding of the relationship between hazard of infection and climate could improve livestock management and animal health. The selected species also have many similarities with soil-transmitted helminths of humans, and insights from their thermal dynamics can be used for public health prevention in endemic areas under climate warming.

## Materials and methods

### Literature search and data analysis

We performed a systematic review of published studies on the effect of temperature and relative humidity on free-living stages of soil-transmitted Trichostrongylidae of mammal herbivores following PRISMA (https://prisma-statement.org/). We limited our analysis to studies from the laboratory at controlled climatic regimes, which allowed us to include extreme conditions that might not be experienced in current natural settings but could represent future situations due to climate change. We screened an initial number of 244 publications and selected a total of 22 papers that met our criteria. Nine species were identified from five genera (*Trichostrongylus, Haemonchus, Ostertagia, Cooperia, and Nematodirus*) (SI.1, Table S1 and Figures S1,S2).

An initial analysis was undertaken to examine if the ecological and phylogenetic diversity of the considered species could explain differences in the traits of their stages free-living in the environment, specifically egg hatching and survival, 1^st^stage larval development, and 3^st^ stage larval survival (Figure 1). ANOVA was used to test the contribution of parasite’s phylogeny, host species, and site of infection, including their interactions with climatic variables, to variation in the observed traits (SI.2, Table S2-S4).

**Figure 1.**
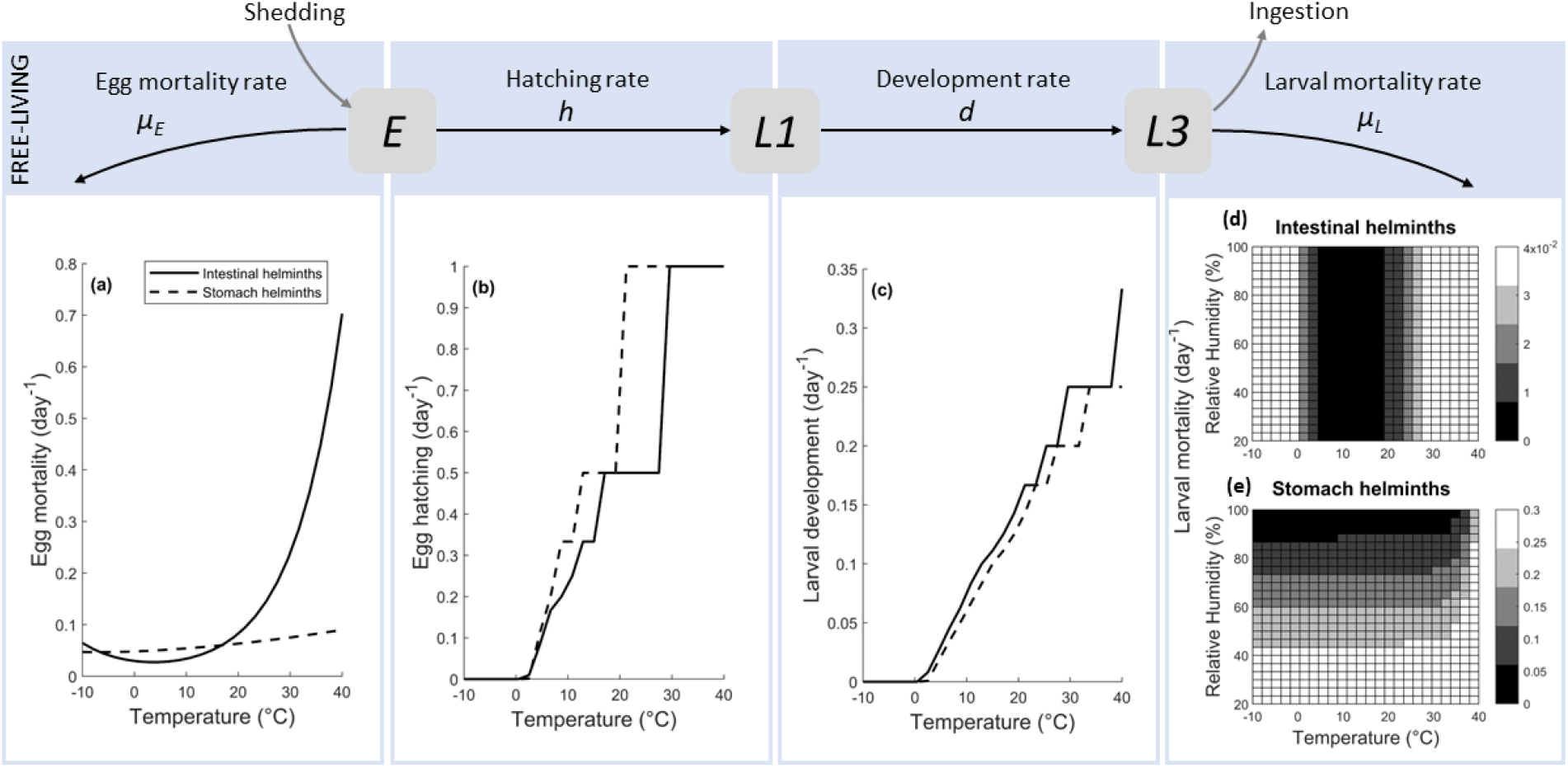
Life cycle of free-living helminths (upper blue panel) and climate dependencies of demographic rates for intestinal (bold lines a-c and d) and stomach (dashed lines a-c and e) helminths from laboratory data.

### Climate-driven model of free-living stages

The free-living component of the life cycle of soil-transmitted helminths can be described by the following set of ordinary differential equations, which captures the dynamics of eggs (*E*), L1-stage larvae (*L*_*1*_), and L3-stage infective larvae (*L*_*3*_) in time with climate-dependent parameters, as follows (Figure 1):

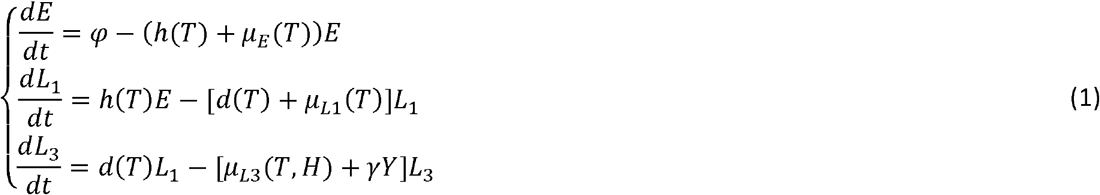

where *t* represents time in days, *φ* is the daily rate of eggs shed in the environment, *h* is the egg hatching rate into L1, μ_*E*_ is the egg mortality rate, *d* is the larval development rate from L1 to L3, μ_*L*1_ and *μ*_*L*3_ are the larval mortality rate of L1 and L3, respectively, *γY* and represents the uptake of infective larvae by the hosts, with and *γ* being *Y* the grazing rate and the host abundance, respectively. The three stages are affected by mean air temperature (*T*, thereafter referred to as temperature, °C) and/or air relative humidity (*H*, thereafter referred to as humidity, %). Eggs and L1 live in the host’s feces and we assumed that feces maintain the minimum humidity required for their viability; in contrast, L3 larvae emerge from the feces to live on the pasture and are under the direct influence of climatic conditions. For simplicity, we assumed that the mortality of L1 in the feces is negligible (*γ*_*L*1_=0) and omitted to model the intermediate larval stage L2, which also lives in the feces and is morphologically similarities to L1.

We expressed the rate of egg hatching as a temperature-dependent process, described by the following Degree-Day (DD) functions (Dobson et al., 2015; Hernandez et al., 2013; Hsu and Levine, 1977):

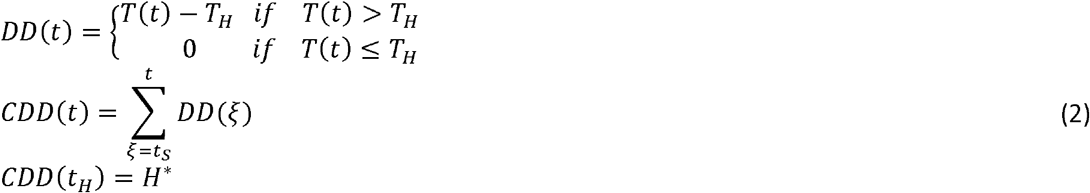

where *t*_*S*_ and *t*_*H*_ (days) are the egg shedding time and the hatching time, respectively, *T*_*H*_ (°C) is the baseline temperature for *DD* accumulation, and *H*\* is the Cumulative *DD (CDD)* threshold needed by eggs to hatch. The thermal-dependent mortality of eggs was modeled as a sum of exponential functions of temperature described by the non-negative parameters *b*_*1E*_, *β*_*1E*_, *b*_*2E*_, and *β*_*2E*_ (Smith, 1990) as:

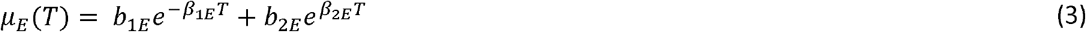

The rate at which larvae L1 develop into infective L3 depends on temperature following a *DD* model similarly to equation 2 for egg hatching (Dobson et al., 2015; Hernandez et al., 2013; Hsu and Levine, 1977):

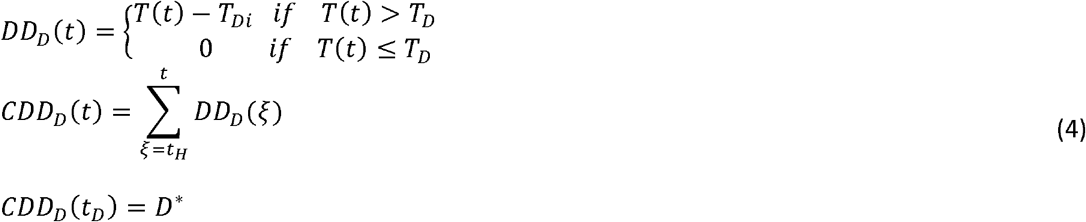

with *t*_*D*_ representing the development time in days, *T*_*D*_ being the baseline temperature for degree day accumulation (°C) and *D*\* being the *CDD* threshold needed by L1 to develop into L3 larvae.

We assumed that the L3 mortality rate depends both on the daily temperature and humidity, with a functional relationship that was modeled as a sum of exponential functions of temperature (Smith, 1990) and as a linear function of humidity and as a linear function of humidity (Dagostin et al., 2023; Mignatti et al., 2016) as:

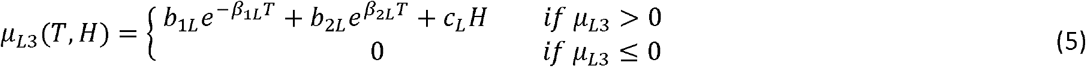

Where the non-negative parameters *b*_*1L*_, *β*_*1L*_, *b*_*2L*_, and *β*_*2*L_ depict the impact of temperature while *c*_L_ describes the effect of humidity.

We used the gathered experimental observations grouped in intestinal and stomach helminths (see SI.2 for details) and calibrated the climate-driven functions of egg hatching and mortality, L1 development and L3 mortality (eqs. 2-5) minimizing the root mean square error (*ERR*) for each group independently as:

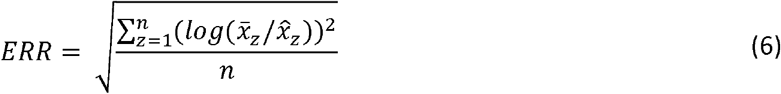

Where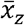and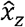represent the observed and estimated, via the described model equations, free-living rates, respectively, which were log-transformed to better compare rates of different order of magnitude, while *n* is the sample size.

Finally, we evaluated the net effect of the different rates on the pool of infective L3 larvae in the environment. Assuming that *T* and *H* vary on a longer time scale than *E, L*_*1*_, *L*_*3*_, we can calculate L3 at quasi-equilibrium for any given *T* and *H* as:

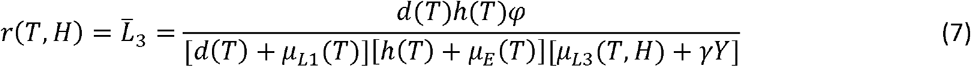

This quantity represents a proxy for the climatic-dependent hazard of infection for any given host exposed to these helminths. To simplify the analysis, the infection hazard was calculated per unit of egg shedding rate (φ=1) while host mortality induced by L3 uptake was considered to be negligible (γ*Y*=0).

To identify the parameters with the strongest effect on the hazard of infection and to evaluate the sensitivity to parameter uncertainty, we implemented the Latin Hypercube Sampling and Partial Rank Correlation (PRC) analysis (Blower and Dowlatabadi, 1994; Marino et al., 2008; McLeod et al., 2006), independently for stomach and intestinal helminths (SI.5).

### Historical and future hazard to helminth infection in Europe

The developed climate-dependent model was then used to examine the climatic responses of eggs, L1, and L3 rates and the resulting hazard of infection intra-annually and throughout Europe, for stomach and intestinal helminths. Using daily mean temperature and relative humidity estimates from the EUROCORDEX (European Coordinated Regional Climate Downscaling Experiment) dataset at a spatial resolution of ∼12.5x12.5^2^ km (https://www.euro-cordex.net/), we considered a total of 44,331 spatial cells, reasonably assuming spatial-independence with no movements of eggs or larvae between cells. We approached this analysis in two phases. First, using a daily time step for each cell of the spatial domain, we assessed the infection hazard and related free-living stage rates, both for stomach and intestinal helminths, in the selected historical period of 20-years (1981-2000) to establish a representative baseline scenario. Second, we evaluated changes in the infection hazard of the two groups in a future 20-year period (2071-2090), under the Representative Concentration Pathway (RCP) 8.5 of “business as usual” high emission scenario (IPCC, 2013). Moreover, we chose the RCP 8.5 projections as representative of extreme, albeit not completely unrealistic scenarios, and we expect that more moderate scenarios would lie in-between the historical and the RCP 8.5 trajectories.

To account for the climatic variability of the European territory, we divided Europe in three main latitudinal areas, which correspond to the main European climatic zones, namely Mediterranean, temperate, and Nordic climates for southern, central and northern Europe, respectively (SI.3, Figure S3).

## Results

### Climatic responses of helminth free-living stages and hazard

The observed rates of egg hatching and mortality, larval L1 development, and L3 mortality were significantly different when the site of infection of the nine Trichostrongylidae species was considered (p<<0.05), while neither the phylogenetic distance between species nor their host species had a significant effect (SI.2, Tables S2-S4). Therefore, we clustered the gathered data in two groups, intestinal (total observations=128) and stomach (total observations=162) helminths, and all the subsequent analyses were carried out independently for each group (SI.4, Table 1, Figure S4,S5). Overall, we found contrasting thermal responses between the two groups. Intestinal helminths showed to have lower baseline temperatures (*T*_*H*_ and *T*_*D*_) and higher degree day (*DD*) cumulated thresholds (*H*^*^ and *D*^*^) both for egg hatching, but not for L1 development, compared to stomach helminths (Figure 1b,c and Table 1). Moreover, the egg mortality rate was more clearly affected by temperature for intestinal than stomach helminths (Figure 1a). Stronger differences were found for L3 mortality, for intestinal helminths the mortality rate was well described by a skewed U-shaped temperature function with a minimum at 10.5°C, while for stomach helminths it was primarily driven by a negative linear relationship with humidity, and an increase in mortality above the temperature extreme of 30°C (Figure 1d,e).

**Table 1.**
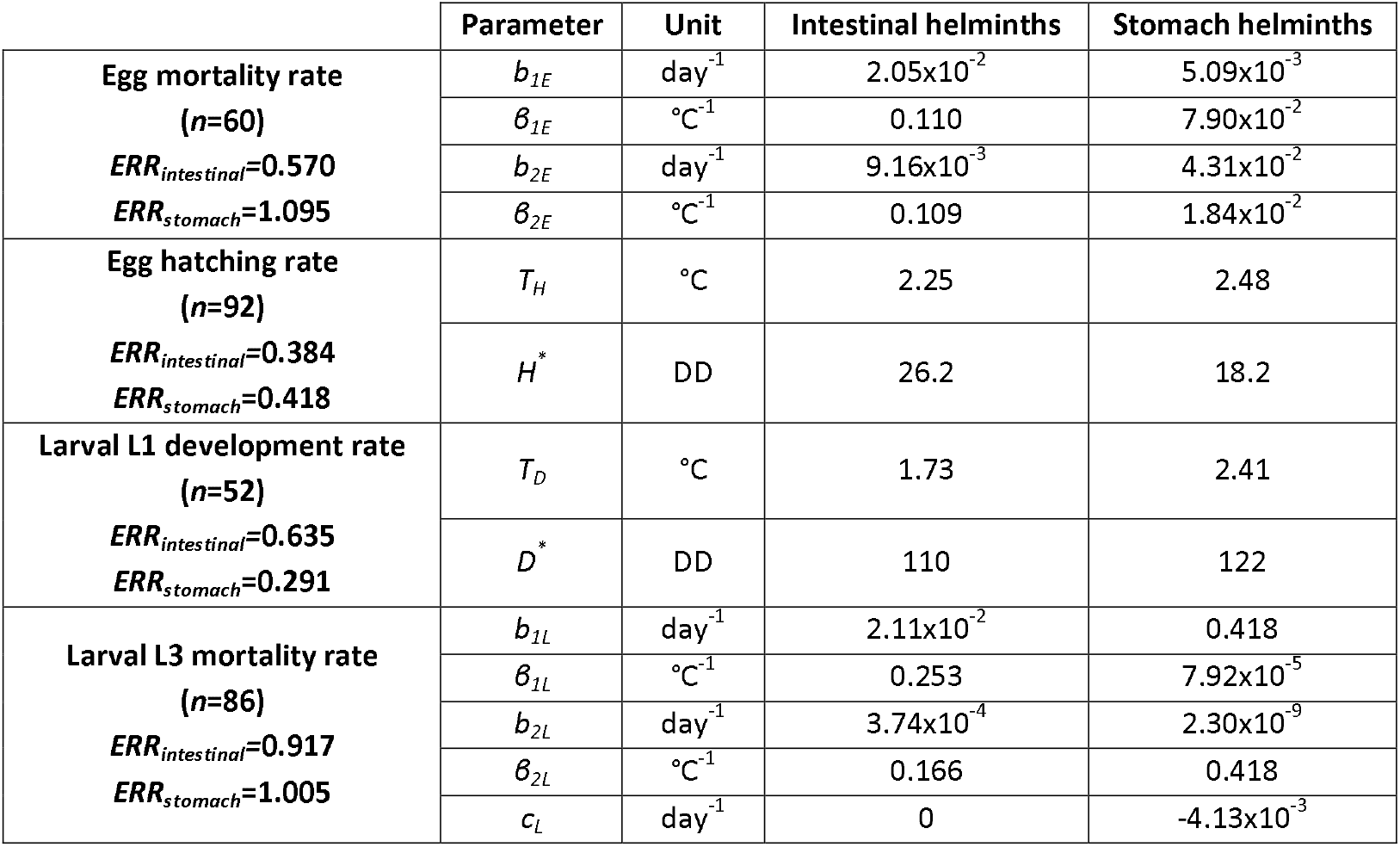
Calibrated parameters (*b*_*1E*_, *β*_*1E*_, *b*_*2E*_, *β*_*2E*_, *T*_*H*_, *H*^*^, *T*_*D*_, *D*^*^, *b*_*1L*_, *β*_*1L*_, *b*_*2L*_, *β*_*2L*_ and *c*_*L*_) for the rates of egg mortality and hatching, larval L1 development and L3 mortality as functions of temperature and relative humidity for intestinal and stomach helminths from laboratory data; n indicates the sample size and ERR represents the Root Mean Square Error described in eq. 6.

We then examined the infection hazard for each helminth group under a range of temperature and humidity conditions, and differences between these two groups were evident (Figure 2). For intestinal helminths, the hazard was strongly affected by temperature peaking at the optimal temperature of 10°C, which corresponds to the highest L3 survival, and dropping to zero below 2°C and above 35°C

**Figure 2.**
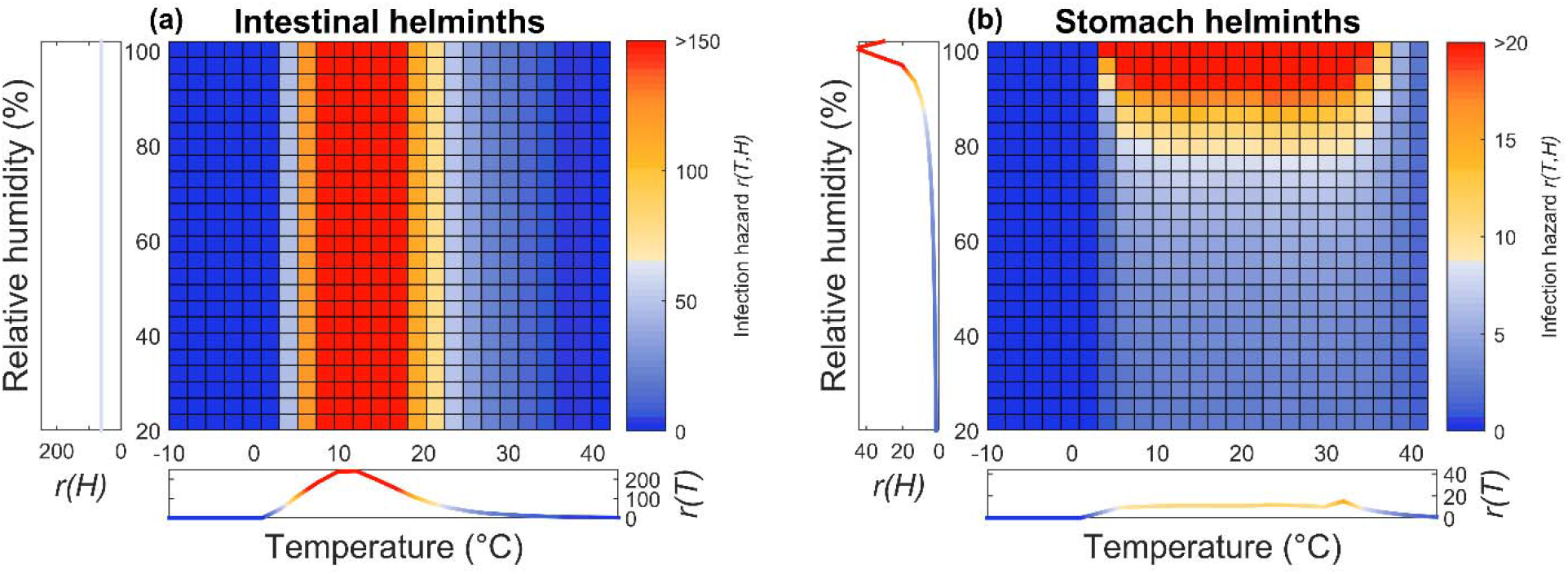
Response of the infection hazard *r(T,H)* of (a) intestinal and (b) stomach helminths under different temperatures (x-axis) and relative humidities (y-axis), estimated from laboratory data. Profiles on the x- and y-axis represent the average infection hazard by temperature *r*(*T*), and by humidity *r(H*), respectively.

because of a decrease in egg hatching and development or high L3 mortality, respectively. Humidity appeared to have no clear effect on the hazard of intestinal helminths (Figure 2a). In contrast, humidity substantially impacted the hazard of stomach helminths in that the greatest hazard was found at above 80% humidity within the 5°C-35°C temperature range (Figure 2b). The partial rank correlation coefficients from the performed sensitivity analysis reinforce our findings by showing that the relative contribution of the different parameters to the infection hazard consistently differs between the two helminth groups (Figure S6).

These findings highlight the contrasting climatic performance of intestine and stomach helminths when exposed to the same temperature and humidity conditions. Moreover, these differences show that the net climatic effect on the infection hazard is determined by the complex balance between different demographic traits and their thermal dependence, namely, how fast eggs can hatch and for how long eggs and infective L3 can remain viable in the environment.

### Historical and future seasonality of helminth hazard

The relationships between climatic variables and demography of free-living helminths, previously identified and trained on laboratory data, were then used to investigate how the infection hazard is affected by seasonality in the three European zones for the historical (1981-2000) period and future (2071-2090) projections under the RCP 8.5 climate change scenario (Figures 3, S7, SI.6).

**Figure 3.**
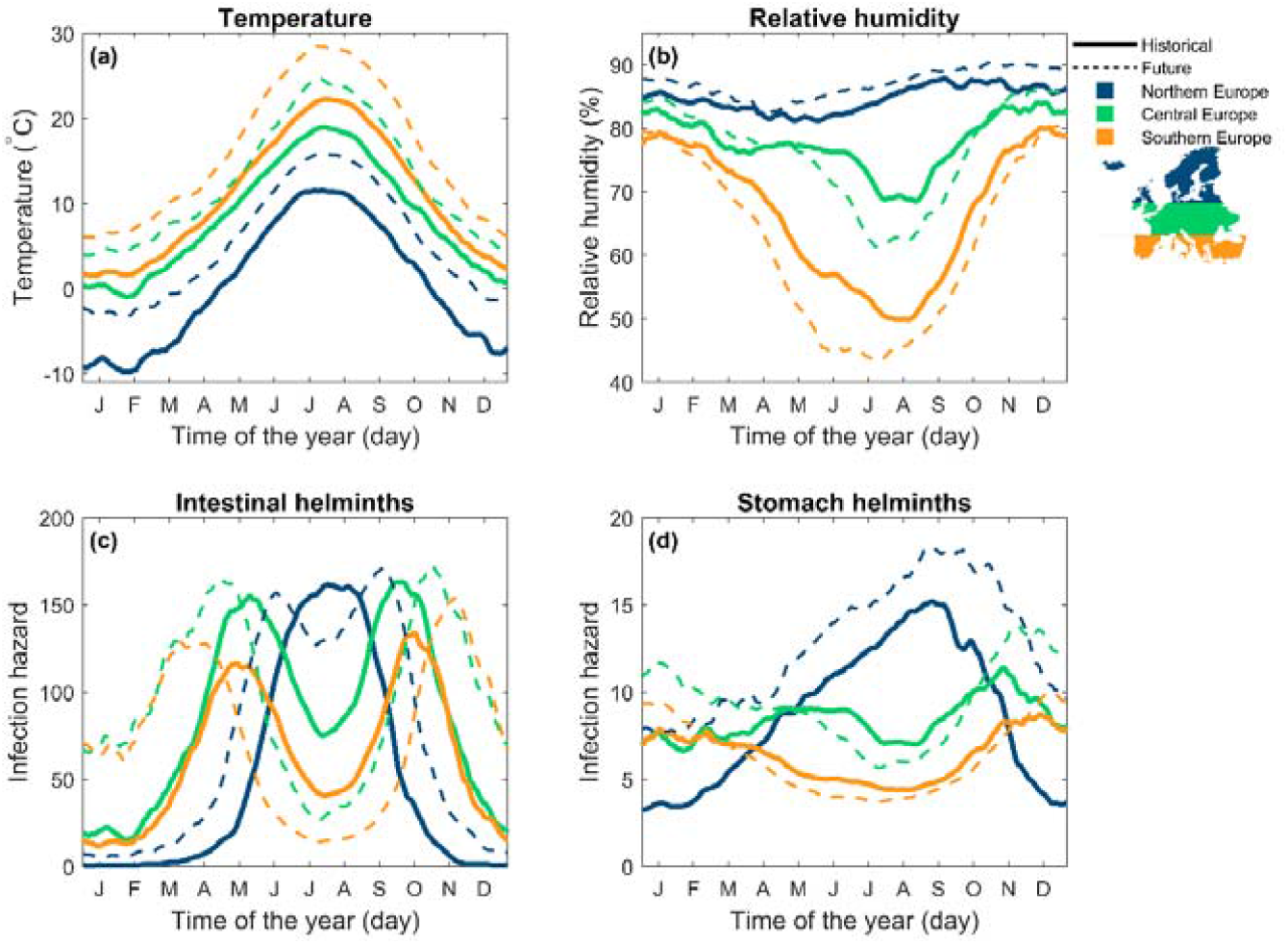
Average historical (1981-2000, bold lines) and future (2071-2090 under RCP 8.5 scenario, dashed lines) seasonality of (a) temperature, (b) relative humidity, and (c,d) infection hazard of (c) intestinal and (d) stomach helminths in northern (blue), central (green), and southern (orange) Europe, plotted with two weeks moving average for visual representation.

During the selected historical period, temperature exhibited the classical annual unimodal profile with a peak in summer in all the three European zones and variation in the absolute values according to their latitudinal gradient (Figure 3a). The seasonal trend of humidity was more variable among the three zones: humidity was above 80% and relatively constant in northern Europe but exhibited a summer minimum more pronounced in southern than central Europe (Figure 3b). Our model simulations indicate that climate seasonality led to contrasting patterns in the hazard of infection between the two helminth groups and among the geographical zones within each group (Figure 3). The hazard of intestinal helminths exhibited a unimodal trend peaking in summer for northern Europe and a biannual pattern with peaks in spring and fall for central and southern Europe (Figure 3c). The striking difference between latitudinal zones appears to be caused by the summer mismatch between the rates of hatching/development and mortality, being the latter greater for eggs and larvae at low latitudes compared to high latitudes (Figure S7). The hazard of stomach helminth infection depicted a different scenario characterized by a late-summer peak in northern Europe, probably facilitated by a generally low L3 mortality coupled with fast egg hatching and L1 development in summer (Figure 3d, S7). In the remaining European zones, the hazard followed the humidity profile, except for winter months when cold temperatures are a restraining factor for hatching and development (Figure 3d). At lower European latitudes, the two helminth groups showed the most compelling seasonal differences, suggesting strong differences in the fulfillment of their ecological requirements.

Projections at the end of the XXI century under the RCP 8.5 scenario indicated that climate change is expected to cause an increase of temperatures, which will maintain the same seasonal trends for the three European zones (Figure 3a). Humidity is also projected to change and exacerbate the preexisting latitudinal differences by causing a more humid northern Europe and drier central and southern zones, particularly in summer months (Figure 3b). Projections suggested that the climatic responses of free-living stages will reflect these seasonal changes by showing an increase in the dissimilarities between the two helminth groups, already highlighted during the historical period. The bimodal seasonality noted for intestinal helminths in the historical period is expected to become more prominent with the shift and increase of the two peaks towards early spring and late fall. We also expect the emergence of a similar bimodal trend in northern Europe and a general increase of the hazard of infection in the winter months in three European zones (Figure 3c). The intensification of this bimodality is mainly caused by a lower egg and larval survival during the summer months (Figure S7). The hazard of stomach helminth infection will drastically increase throughout the year in northern Europe. At lower latitudes, the coupling of drier and warmer conditions from spring to fall is expected to negatively affect the hazard, while warmer and slightly wetter winters will favor infections (Figure 3).

In summary, climate warming is expected to increase the intensity and timing of the hazard of infection of both helminth groups on the pasture in the fall-winter months but will have an opposite effect in the summer months; this change will also be associated with a temporal shift and increase of the peak.

### Historical and future spatial distribution of helminth hazard

We examined the spatial distribution of the helminth hazard during the selected historical period (1981-2000) and the expected percentage changes in the future (2071-2090) under the RCP 8.5 climate change scenario. (Figure 4, S8, S9, SI.7).

**Figure 4.**
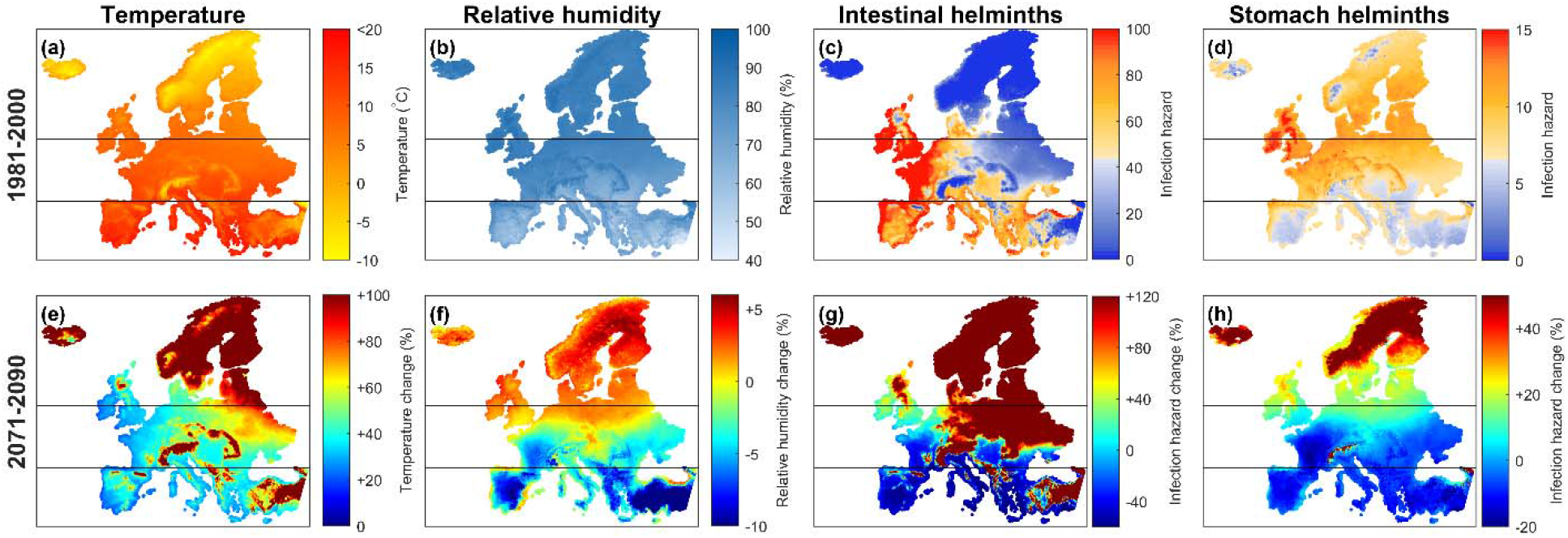
Average historical (1981-2000) (a) temperature, (b) relative humidity, hazard of infection of (c) intestinal and (d) stomach helminths and future (2071-2090) expected changes (e-h) under the RCP 8.5 climate change scenario. Horizontal lines separate northern, central and southern zones.

The historical spatial distribution of the average annual temperature and humidity confirms the well-characterized latitudinal trends (Figure 4a,b). These geographical differences were found to have contrasting effects on the spatial distribution of the demographic rates withintestinal helminths exhibiting a minimum hazard of infection in the north-eastern part of the continent, which on the contrary, represented an infection hotspot with high hazard for stomach helminths (Figures 4c,d, S8).

The projected changes of temperature and humidity at the end of the XXI century will not occur uniformly throughout Europe (Figure 4e,f). Southern and northern Europe are expected to experience a drastic warming, up to +6 C, with a decrease of humidity in the former (-10%) and an increase in the latter (+5%) zone. In central Europe, temperature is projected to increase by +4 C, but humidity should remain mostly unchanged. Consequently, Scandinavian countries are expected to experience a greater hazard of infection for both helminth groups (increase of 100% for intestinal helminths and 55% for stomach helminths) due to a fast egg hatching and L1 development paired with high egg and L3 survival (Figures 4g,h, S9). On the contrary, the substantial warmer temperatures and drier conditions in the Mediterranean countries are projected to reduce the hazard of both helminth groups (decrease of 60% for intestinal helminths and 20% for stomach helminths) due to a drastic increase of egg and L3 mortality and (Figures 4g,h, S9). Additionally, our projections highlight dissimilarities between the two groups for high altitude areas, such as Pyrenees, Alps, Pontic mountains, where we expect a higher increase in the hazard of intestinal helminth infections compared to stomach helminths (Figure 4g,h). Overall, in addition to identifying a spatial shift towards northern latitudes of the hazard of infection and associated decrease at lower latitude for both helminths, our maps confirm the strong regional variation in the climate-hazard relationship.

### Spatiotemporal changes of helminth co-occurrence

The co-occurrence of intestinal and stomach helminths was evaluated to examine the degree of spatial overlapping and to determine whether areas of co-circulation may expand in the future (SI.8, Figure S10).

In central Europe, the co-occurrence of intestinal and stomach helminths (82% of spatial cells) is facilitated by climatic conditions favorable for both groups, particularly intermediate temperatures and high humidity for intestinal and stomach helminths, respectively. Southern Europe generates climatic areas where single circulation of intestinal species predominates (57%), whereas single circulation of stomach species appears more common in northern Europe (65%) (Figure 5a,b). Unfavorable areas to both helminth groups are only represented by 4% of the entire European territory and are mainly located in the Alps, Scandinavian mountains, and Iceland. When we examined the average infection hazard across the latitudinal gradient, results confirmed that the co-circulation of both groups in the environment predominates, except for very high latitudes where single infection by stomach helminths appears to prevail (Figure 5c,d).

**Figure 5.**
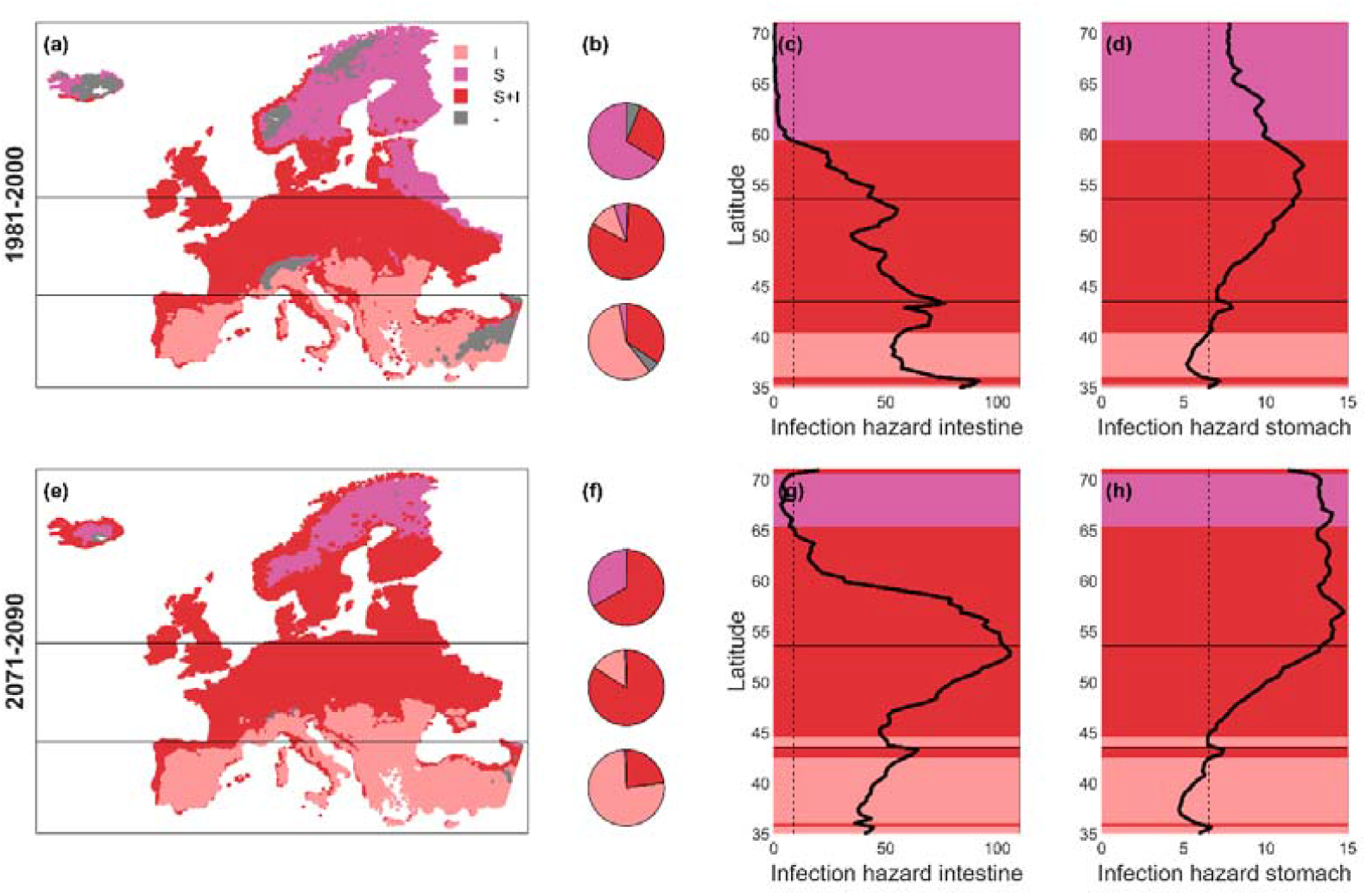
Historical (1981-2000) (a-d) and future (2071-2090) (e-h) single occurrence of intestinal helminths (I, pink), single occurrence of stomach helminths (S, purple), co-occurrence of both helminth groups (S+I, red), and no occurrence of both groups (-, grey) in Europe. Spatial distribution (a,e), composition of spatial cells (b,f) in northern Europe (top pie), central Europe (mid pie), southern Europe (bottom pie) based to their occurrence type, and average infection hazard of intestinal (c,g) and stomach (d,h) helminths. Black bold lines (c,d,g,h) represent the average infection hazard across the European latitudes, the vertical dashed black line is the threshold r and the color in the background represents the average occurrence type at each latitude.

Future projections indicate an expansion of the distribution of small intestine helminths in the Mediterranean regions (+19%) and co-circulation of both groups in northern Europe (+39%, Figure 6e,f). Unfavorable areas to helminth infection are expected to decrease in the future (from 4% to less than 1%). Overall, our predictions suggest that for mid-high latitudes the probability of co-occurrence of stomach and intestinal helminths in the environment is expected to be higher given that the relative hazard of infection of both groups will substantially increase.

## Discussion

We developed a climate-driven mechanistic model to examine the demographic responses of trichostrongylid helminths, aggregated in intestinal and stomach groups, and their hazard of infection. This framework was then applied at the European continental scale to evaluate changes in the seasonal and spatial distribution of the two groups during historical and projected climatic scenarios. Our predictions suggest that under the high emission scenario RCP 8.5 climate will severely modify the seasonal transmission and the spatial distribution of the two groups according to their distinct ecological requirements and climatic tolerance along the latitudinal and altitudinal gradients. A drastic increase of the infection hazard at mid-high latitudes will likely increase the co-circulation of stomach and intestine helminth species with critical consequences for animal health in the future.

We found clear differences in the response of helminths to temperature and humidity based on their site of infection in the mammal host. Multiple ecological factors could contribute to explaining this contrasting trend. One possibility could be related to the intrinsic differences in the life history properties between the two groups. For example, the body size of adult female is bigger in stomach than intestine helminths, irrespective of their mammal host, a pattern that is partially reflected in the volume of their eggs (Table S1). Individual body size is a fundamental characteristic that influences the thermal sensitivity of an organism, especially for ectotherms, where larger sizes are commonly associated with higher temperature tolerance and fitness (Kingsolver and Huey, 2008). It is indeed possible that the bigger stomach helminths produce eggs, and likely infective larvae, of better quality and larger size that can thrive under high temperatures. Differences in the immune response between the two groups, and likely affecting female fecundity including egg quantity and quality, could also contribute to the observed patterns. Previous studies have suggested that host immunity can constrain the fitness, in terms of body size and/or egg abundance in uterus, of adult helminth females and consequently the hatching potential of their eggs (Hein et al., 2010; Lambert et al., 2015; Viney and Cable, 2011; Wakelin, 1987). For example, a negative relation between the host immune response and both female length and number of eggs in uterus was found for the stomach helminth O. circumcincta in sheep (Stear et al., 1995) and the intestinal T. retortaeformis in rabbits (Cattadori et al., 2005, 2019, 2014; Chylinski et al., 2009). Given the complex relation between host immunity and helminth fitness, we cannot exclude that the immune response could affect the way helminths respond to different climatic conditions.

Our model projections suggest that there are significant differences in the seasonal and spatial trends of the hazard of infection both between and within helminth groups throughout Europe. The emergence of a bimodal seasonal hazard with a minimum in summer for intestinal helminths is determined by the overshoot of the thermal optimum during extreme hot periods, a profile described for the historical time in central-southern Europe and likely to be exacerbated in future warmer conditions. Importantly, a similar bimodal pattern is expected for the northern regions in the long-term as they are projected to experience extreme hot temperatures. These findings are in agreement with climate change predictions from theoretical models (Altizer et al., 2013; Molnár et al., 2013a), and frameworks applied to trichostrongylid infections under different climate change scenarios (Rose et al., 2016, 2015). For stomach helminths, whose spatiotemporal dynamics are mainly affected by humidity, central-northern Europe could be potentially expose to a higher infection hazard due to relative warm and humid summers within the tolerance range of these helminth species. Our simulations indicate that stomach helminths are distributed across Europe following an increasing south-north gradient, which is the opposite for the intestinal helminths. Noteworthy, the infection hazard of both helminth groups is expected to expand towards the northern latitudes and contract in the southern territories. These climate-driven spatial shifts are in agreement with projected distributions of helminths of wild (e.g. Dall sheep, muskoxen) and farm (e.g. sheep) animals in northern Europe and Canada, where a more permissive climate is already contributing to expand the spatial ranges of the local parasite species (Jenkins et al., 2006; Kutz et al., 2009, 2005b; Rose et al., 2016). Similarly, the warming of the artic tundra has been shown to likely allow for the completion of the parasite life cycles facilitating the establishment and expansion towards northern areas (Kafle et al., 2020; Kutz et al., 2013). Anticipating the spatiotemporal shifts of the hazard of helminth infection is particularly important from an applied perspective. At present, helminth treatment represents 20% of the total economic burden to the livestock industry in Europe (Charlier et al., 2020) and an alarming positive trend is expected to continue in the future due to the spread of anthelminthic resistance (Geerts and Gryseels, 2001; Rose Vineer et al., 2020) and/or the lack of regular prophylactic. Redesigning the timing and the area of interest of these treatment strategies could reduce the spread of anthelminthic resistance and contribute to a more effective control of helminth infections.

By investigating the response of closely related species exposed to the same climatic drivers, our work provides a more comprehensive understanding of the climate change impact on the co-circulation and potential coinfection of multiple helminth species. Model projections identify the European mid-high latitudes as future critical areas for the co-circulation of intestinal and stomach helminths, with an expected increase of the hazard by +100% and +30%, respectively. Moreover, future climate changes might increase the likelihood of synchronous seasonal peaks of infection hazard to both groups. This spatial and seasonal overlapping could facilitate coinfection events, given the exposure to a competent host, and increase disease severity and transmission with consequences for species co-endemicity and infection hotspots. Recently, we showed that while under the same climatic conditions coinfected rabbits shed more *T. retortaeformis* and *G. strigosum* eggs that survive for longer in the environment, compared to single-infected hosts (Dagostin et al., 2023), supporting the important role that co-infected individuals could play in disease transmission under climate change.

Our results bring a new perspective to the study of climate and infectious diseases by addressing the role of humidity as climatic driver of helminth transmission. Indeed, despite the well-recognized humidity requirements for helminth life cycles (Beveridge et al., 1989; Pandey et al., 1993; Prasad, 1959) this variable is usually neglected (but see (Dagostin et al., 2023; Maya et al., 2010)) in favor of temperature, which is more frequently recorded in the field and easy to manipulate in the laboratory. We show that humidity is a key environmental variable for understanding the climate-driven dynamics of stomach helminth free-living stages and important for generating heterogeneities in the demographic responses of both intestinal and stomach group. Moreover, the inclusion of humidity provides a more accurate understanding of the relationship between climate and hazard of infection, including a more rigorous projection under future climate change.

Ultimately, our work shows that the ecological requirements of different soil-transmitted helminth species modulate whether and how climate change will modify the magnitude, seasonality and distribution of the infection hazard and the co-circulation of multiple helminth species over time and space.

