## Supplementary material for "Helminth ecological requirements shape the impact of climate change on the hazard of infection": SI

**Title**

**Supplementary Information**

**SI.1 Literature analysis**

The Preferred Reporting Items for Systematic Reviews and Meta-Analyses (PRISMA) was used to identify available publications on the effect of climate on the soil-transmitted free-living stages of the most common Trichostrongylidae of mammal herbivores from laboratory experiments. We used the following keywords in the literature search:

( ( ( "Temperature"  OR  "Humidity" )  AND  ( "Trichostrongylus"  OR  "Ostertagia"  OR  "Teladorsagia"  OR  "Graphidium"  OR  "Haemonchus" )  AND  ( "egg"  OR  "larvae" )  AND  ( "survival"  OR  "development

OR "hatching" ) ) ).

We screened an initial number of 244 published records, we excluded 201 records as they did not meet the our criteria and we selected a total of 22 published papers (Andersen and Levine, 1968; Beveridge et al., 1989; Boag and Thomas, 1985; Ellenby, 1968; Gibson, 1981; Hernandez et al., 2013; Jehan and Gupta, 1974; Lambert et al., 2015; Mirzayans, 1969; Pandey, 1972a, 1972b; Pandey et al., 1993, 1989; Prasad, 1959; Rose, 1963, 1961; Rose and Small, 1984; Salih and Grainger, 1982; Silverman and Campbell, 1959; Sonibare et al., 2011; Wharton, 1982; Young et al., 1980) (Figure S1,S2).

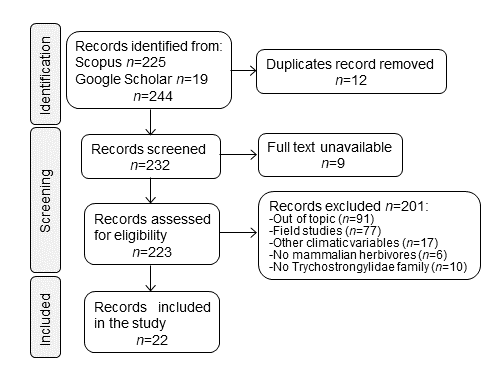

**Figure S1.** PRISMA literature analysis of the climate effects on the free-living stages of parasitic Trichostrongylidae.

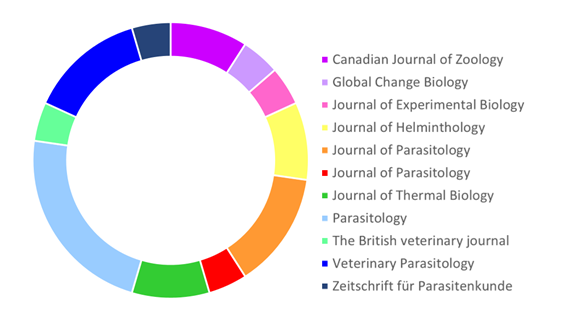

**Figure S2.** Journal publications of the selected papers of performed PRISMA literature analysis.

From the selected 22 published papers we compiled data on nine species that infect the intestine or the stomach of different hosts (see Table S1).

Table S1 Summary of epidemiological, morphological, and physiological characteristics of the considered helminths. Egg volume is calculated assuming that the egg shape can be approximated to an oval.

| **Species** | **Main host** | **Infection Site** | **Adult length (mm)** | **Egg size (μm)** | **Egg volume**  **(μm^3^)** | **Phylogenetic distance ^α^** |
| --- | --- | --- | --- | --- | --- | --- |
| *T. rugatus^Φ^* | Sheep | Small intestine | M=4-6 F=5-8 | 90x40 | 75,000 | 0.0933 |
| *T. colubriformis*^𝛽^ | Sheep | Small intestine | M=4-5 F=5-7 | 85x40 | 71,000 | 0.0925 |
| *T. vitrinus*^𝛽^ | Sheep | Small intestine | M=4-6 F=5-8 | 100x45 | 106,000 | 0.0900 |
| *T. retortaeformis^ɣ^* | Rabbit | Small intestine | M=5-8, F=8-9 | 90x40 | 75,000 | 0.0842 |
| *T. axei*^𝛽^ | Cattle | Stomach | M=3-4, F=4-5 | 85x45 | 90,000 | 0.0858 |
| *G. strigosum^δ^* | Rabbit | Stomach | M=12, F=16 | 95x50 | 120,000 | 0.0825 |
| *O. ostertagi^θ^* | Cattle | Stomach | M=6-7, F=8-11 | 86x70 | 220,000 | 0.141 |
| *H. contortus*^𝛽^ | Sheep | Stomach | M=10-20, F=18-30 | 70x44 | 71,000 | 0.103 |
| *T./O. circumcincta*^𝛽^ | Sheep | Stomach | M=5-10, F=6-12 | 90x50 | 120,000 | 0.0800 |

^α^Chilton et al. 2001 (Chilton et al., 2001)

^𝛽^Taylor et al. 2015 (Taylor et al., 2015)

*^δ^*Massoni et al.2011 (Massoni et al., 2011)

*^ɣ^*Audebert et al. 2022 (Audebert et al., 2002)

^Φ^Roeber et al. 2013 (Roeber et al., 2013)

*^θ^*Lichtenfels and Hoeberg 1993 (Lichtenfels and Hoberg, 1993)

**SI.2 Statistical analysis**

From the selected papers, we estimated the median time in days of egg hatching, L1 development, and L3 survival for each climatic condition that was reported, and then computed the respective rates as their reciprocal in days^-1^. Egg mortality rate was estimated from the egg hatching rate *h* and from the percentage of unhatched eggs at the end of the experiment (*M*), with $\mu_{E}=\frac{Mh}{(1-M)}$.

Given the ecological and phylogenetic diversity of the considered parasite species (Table S1), in addition to climate, we examined if the demographic rates of interest were significantly affected by the site of infection of adult parasites in the host (we considered the most common host species they infect), or the phylogenetic relatedness between parasite species using ANalysis Of VAriance (ANOVA, significance level α=0.05) (Table S2-S4). For each of the nine considered species, we estimated the phylogenetic distance to the most recent common ancestor based on Chilton et al. (2001) (Chilton et al., 2001). Categorical variables were used to describe the common host of each helminth and the site of infection of adult parasites within the host (Table S1).

Infection site, climate and/or their interaction significantly explained the observed variance of egg hatching, L1 development, and L3 mortality rates (Table S2). The phylogenetic distance between species and the common host were not significant, except in the two-way interaction with temperature and only for L3 mortality rate (Table S3). Therefore, we clustered the data on the host’s site of infection, ‘Stomach’ or ‘Intestine’, and all the subsequent analyses were performed on each group independently.

**Table S2.** ANOVA of egg mortality, egg hatching, L1 development, and L3 mortality rates (in logarithmic scale) according to climatic variables and infection site.

| **Egg mortality rate** | ***Sum Sq.*** | ***F*** | ***p*** |
| --- | --- | --- | --- |
| Temperature | 11.33 | 11.26 | 0.0014 |
| Infection site | 4.06 | 4.04 | 0.0493 |
| Temperature:Infection site | 5.40 | 5.37 | 0.0242 |
| Error | 56.36 |  |  |
| **Egg hatching rate** | ***Sum Sq.*** | ***F*** | ***p*** |
| Temperature | 59.9 | 329 | <1x10^-15^ |
| Infection site | 0.842 | 4.29 | 0.0342 |
| Temperature:Infection site | 0.364 | 2 | 0.160 |
| Error | 16.0 |  |  |
| **Larval L1 development rate** | ***Sum Sq.*** | ***F*** | ***p*** |
| Temperature | 40.8 | 157 | <1x10^-15^ |
| Infection site | 0.575 | 2.21 | 0.142 |
| Temperature:Infection site | 1.18 | 4.55 | 0.0366 |
| Error | 16.9 |  |  |
| **Larval L3 mortality rate** | ***Sum Sq.*** | ***F*** | ***p*** |
| Temperature | 38.7 | 11.8 | 0.0009 |
| Humidity | 19.6 | 6.03 | 0.0162 |
| Infection site | 15.3 | 4.71 | 0.033 |
| Temperature:Infection site | 43.1 | 13.2 | 0.0005 |
| Humidity:Infection site | 57.1 | 17.6 | 0.0001 |
| Error | 263 |  |  |

**Table S3.** ANOVA of egg mortality, egg hatching, egg mortality, L1 development, and L3 mortality rates (in logarithmic scale) according to climatic variables and phylogenetic distance.

| **Egg mortality rate** | ***Sum Sq.*** | ***F*** | ***p*** |
| --- | --- | --- | --- |
| Temperature | 0 | 0 | 0.999 |
| Phylogenetic distance | 1.448 | 1.35 | 0.441 |
| Temperature:Phylogenetic distance | 0.584 | 0.55 | 0.261 |
| Error | 59.85 |  |  |
| **Egg hatching rate** | ***Sum Sq.*** | ***F*** | ***p*** |
| Temperature | 4.69 | 24.9 | <1x10^-5^ |
| Phylogenetic distance | 0.113 | 0.60 | 0.441 |
| Temperature:Phylogenetic distance | 0.293 | 1.55 | 0.261 |
| Error | 16.6 |  |  |
| **Larval L1 development rate** | ***Sum Sq.*** | ***F*** | ***p*** |
| Temperature | 2.55 | 9.19 | 0.0035 |
| Phylogenetic distance | 0.0146 | 0.05 | 0.819 |
| Temperature:Phylogenetic distance | 0.119 | 0.43 | 0.515 |
| Error | 18.04 |  |  |
| **Larval L3 mortality rate** | ***Sum Sq.*** | ***F*** | ***p*** |
| Temperature | 17.5 | 4.32 | 0.041 |
| Humidity | 5.36 | 1.32 | 0.254 |
| Phylogenetic distance | 1.88 | 0.47 | 0.497 |
| Temperature:Phylogenetic distance | 29.5 | 7.27 | 0.0085 |
| Humidity:Phylogenetic distance | 7.05 | 1.74 | 0.191 |
| Error | 328 |  |  |

**Table S4.** ANOVA of egg mortality, egg hatching, L1 development, and L3 mortality rates (in logarithmic scale) according to climatic variables and main host.

| **Egg mortality rate** | ***Sum Sq.*** | ***F*** | ***p*** |
| --- | --- | --- | --- |
| Temperature | 15.67 | 17.75 | <1x10^-3^ |
| Host | 1.73 | 0.98 | 0.382 |
| Temperature:Host | 0.317 | 0.18 | 0.836 |
| Error | 47.67 |  |  |
| **Egg hatching rate** | ***Sum Sq.*** | ***F*** | ***p*** |
| Temperature | 49.1 | 262 | <1x10^-16^ |
| Host | 0.554 | 1.48 | 0.233 |
| Temperature:Host | 0.879 | 2.35 | 0.102 |
| Error | 16.09 |  |  |
| **Larval L1 development rate** | ***Sum Sq.*** | ***F*** | ***p*** |
| Temperature | 33.23 | 155.2 | <1x10^-16^ |
| Host | 0.132 | 0.31 | 0.736 |
| Temperature:Host | 0.312 | 0.73 | 0.487 |
| Error | 13.48 |  |  |
| **Larval L3 mortality rate*** | ***Sum Sq.*** | ***F*** | ***p*** |
| Temperature | 30.73 | 6.85 | 0.0105 |
| Humidity | 3.079 | 0.69 | 0.4097 |
| Host | 0.959 | 0.11 | 0.8987 |
| Error | 368 |  |  |

*Interaction analysis could not be performed because of the limited sample size

**SI.3 European climatic zones**

We divided Europe in three main latitudinal areas: i) southern Europe, represented by spatial cells below 43° 27' 36" of latitude, ii) central Europe, from 43° 27' 36" to 53° 32' 60" of latitude, and iii) northern Europe, at latitudes greater than 53° 32' 60" (Vilà et al., 2021). These three regions correspond to the main climatic zones of Europe, namely Mediterranean, temperate, and Nordic climates for southern, central and northern Europe, respectively. We reported in Figure S3 the main European climatic zones identified by the European Environmental Agency (EEA, <https://www.eea.europa.eu/data-and-maps/figures/climate>) and the simplified zoning used in this study based on latitudinal bands.

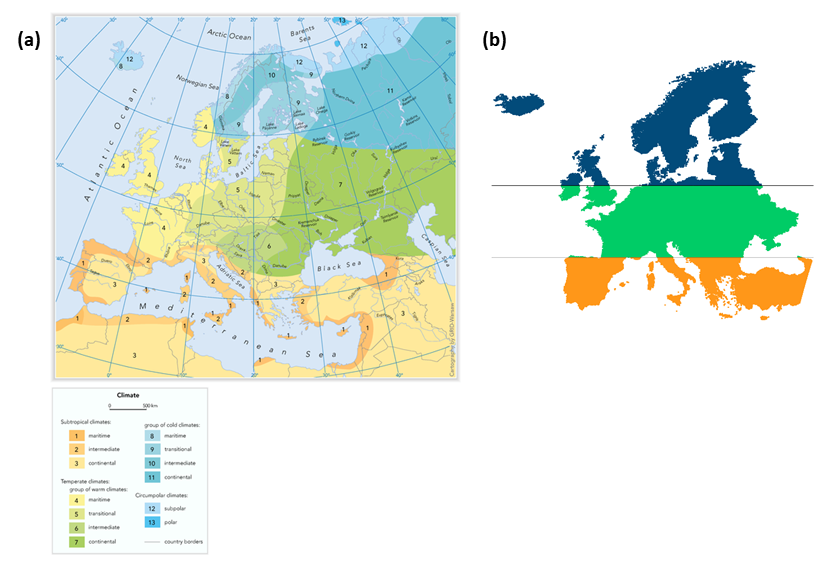

**Figure S3.** European climatic zones (a) identified by the European Environmental Agency and (b) used in this study.

**SI.4 Model calibration**

First, using laboratory data we transformed the observed time for egg survival, egg hatching, larval development and survival into rates and calibrated our model parameters, minimizing the error function described in the main text. Then, we estimated egg hatching time *t_H_*, L1 development time *t_D_* and L3 survival time *t_S_* at different climatic conditions for intestinal and stomach helminths and compared them with the observed data (Figure S4,S5). Overall, our climate-driven model well simulated the observed climatic responses of eggs and L1 larvae. Egg survival of intestinal helminths is well described by a temperature-dependent unimodal curve with a maximum at 4 ̊C (Figure S4a). Both egg hatching and larva development time decrease with temperature and this relationship is well explained by the implemented DD model (Figure S4c-f). There is a weak relationship between egg survival for stomach helminths and temperature confirmed by model fitting (*p*>>0.05, Table S2-S4, Figure S4b). The survival of L3 larvae was well described by a unimodal function that peaked at 10.5 ̊C for intestinal helminths (Figure S5a). The effect of humidity on L3 survival time was almost null for intestinal helminths but strong for stomach helminths, indicating high mortalities at low humidity (Figure S5b).

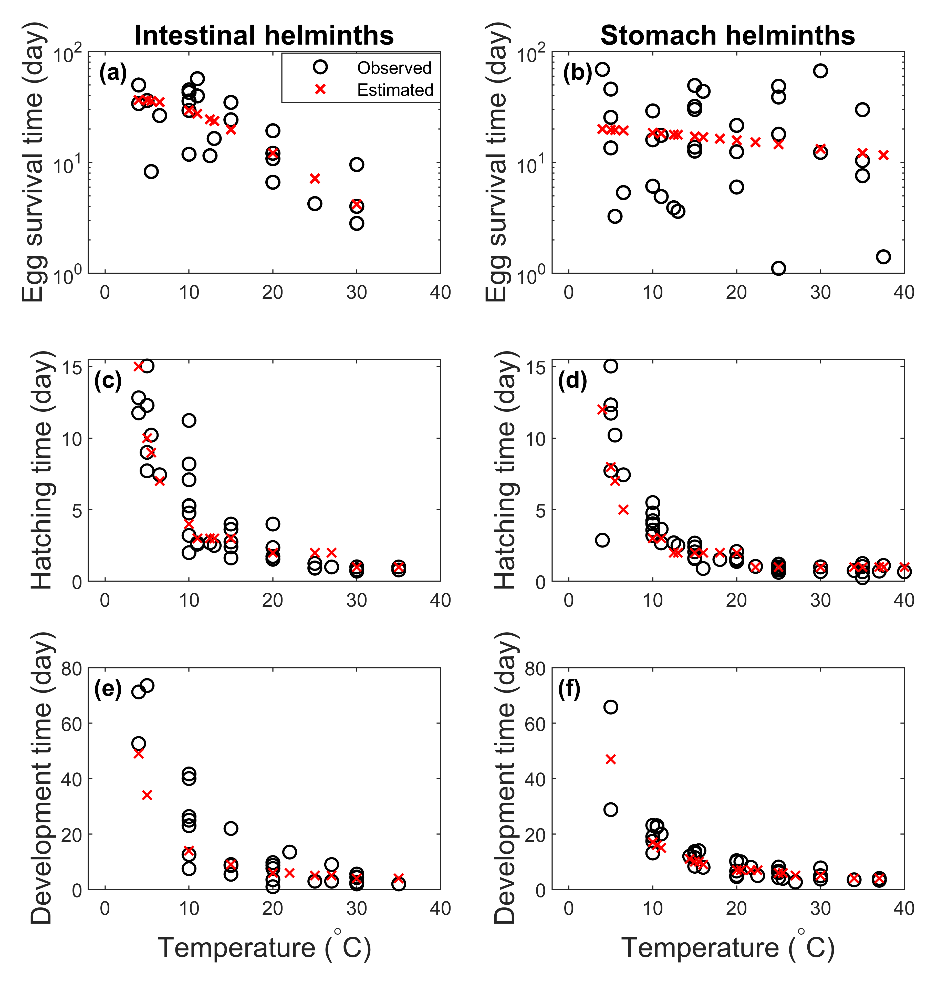

**Figure S4.** Trends of egg survival (a,b), egg hatching (c,d), and L1 development time (e-f) at different temperatures for intestinal and stomach (b,d,f) helminths. Observed (black circles) and simulated (red crosses) values are reported. Note that the y-axis for the egg survival time is in log scale (a,b).

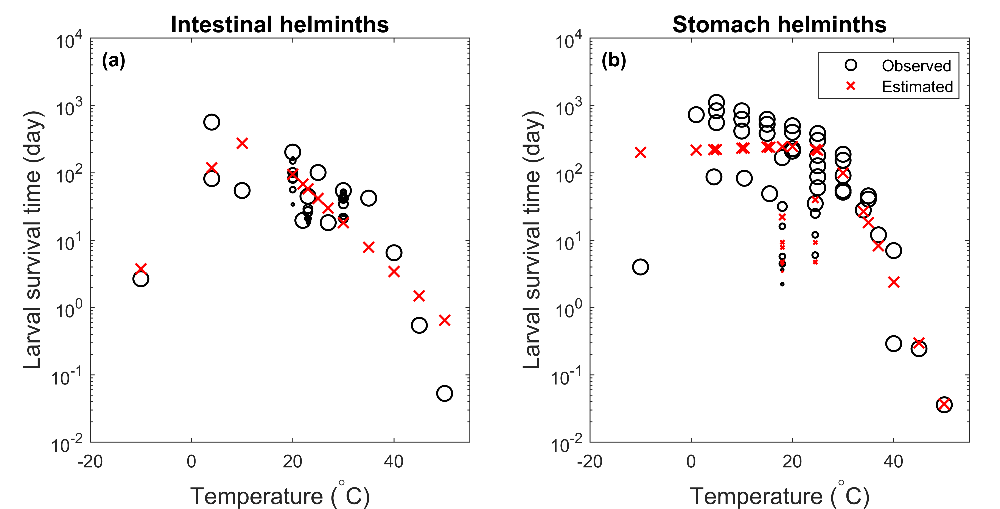

**Figure S5.** Trends of L3 survival time (in log scale) by temperature and at different humidity *H* (circle and cross size) for (a) intestinal and (b) stomach helminths. Observed (black circles) and simulated (red crosses) values are reported. The dimension of the circles and the crosses is proportional to humidity conditions.

**SI.5 Sensitivity analysis on the hazard of infection**

We performed the Latin Hypercube Sampling and Partial Rank Correlation (PRC) analysis (Blower and Dowlatabadi, 1994; Marino et al., 2008; McLeod et al., 2006), independently for stomach and intestinal helminths. This sensitivity analysis was carried out by: i) drawing 100 values for each of the 13 model parameters without replacement from a uniform distribution with extremes in the range of +/- 20% of the optimal parameter values reported in Table 1 of the main text (Gumel et al., 2021); ii) for each row of the 100x13 matrix of parameter values estimated at point *i*, calculating the average infection hazard in the temperature range [-10 °C, +40°C] and in the humidity range [20%, 100%]; and then iii) calculating the PRCs from the parameter matrix and the average infection hazard obtained at each iteration to assess the contribution of uncertainty and variability of individual parameters to uncertainty and variability in the infection hazard. Parameters that show PRC coefficients close to -1 or 1 are expected to have the strongest effect, negative or positive respectively, on the helminth infection hazard (Figure S6).

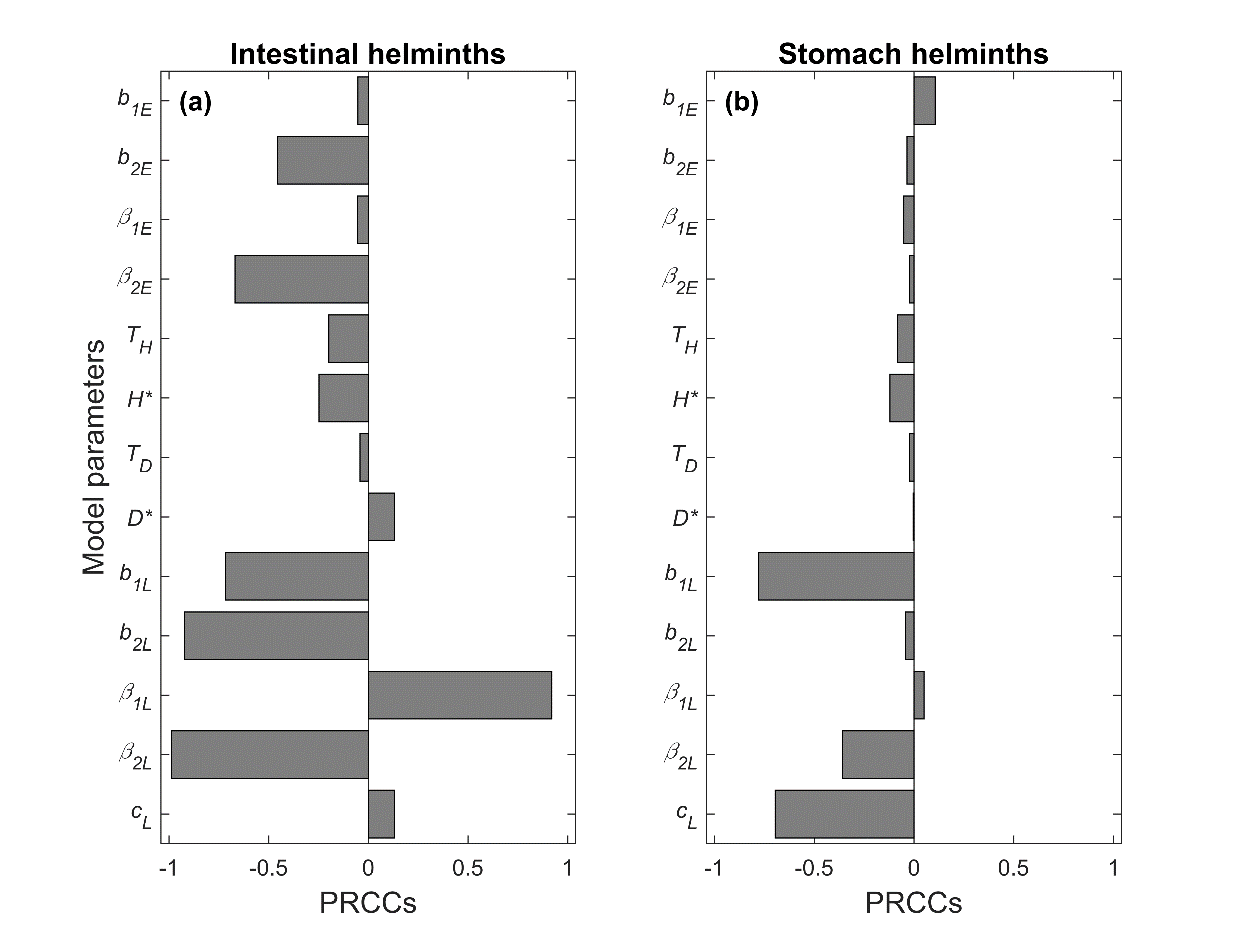

**Figure S6.** Partial rank correlation coefficients (PRCCs) depicting the impact of model parameters on the infection hazard of (a) intestinal helminths and (b) stomach helminths. The baseline values of model parameters are reported in Table 1.

The hazard of intestinal infections is mostly affected by temperature-dependent parameters that describe egg hatching and mortality together with larval survival. Conversely, infection hazard of stomach helminths has the highest contribution from the larval mortality parameters, in particular the humidity-dependent *c_L_* (Figure S6).

**SI.6 Seasonal patterns of demographic rates**

We investigated how the rates of intestinal and stomach helminths were affected by seasonality in the three European zones for the historical (1981-2000) period and future (2071-2090) projections under the RCP 8.5 climate change scenario.

In the considered historical period, egg mortality followed the seasonal temperature profile but in northern Europe, where mortality was high during the winter-fall months and very low in summer, especially for the intestinal group more sensitive to temperature. Egg hatching and L1 development showed similar seasonal trends for both helminth groups and were characterized by a summer peak and higher values in southern Europe followed by the central and the northern zone (Figure S7a-c,e-g). The seasonality of L3 mortality was estimated to be substantially different between the two groups. For intestinal helminths, the highest mortality was concentrated in the winter-fall months, although a much smaller peak was noted in summer for southern and central Europe where high temperatures can further contribute to L3 death. In contrast, L3 mortality of stomach helminths was highest during summer, particularly in southern Europe where dry conditions can increase the mortality of these helminth species, which seem to prefer wet conditions (Figure S7d,h).

In the far future, our simulations suggest that temperature warming will accelerate egg hatching and L1 development for both stomach and intestinal helminths across Europe, but changes will not be consistent for egg and L3 mortality (Figure S7a-d). For intestinal helminths, both rates will drastically decrease in winter-fall because of the less severe temperatures but will increase in summer as temperature is predicted to increase throughout Europe. The mortality rates of egg and L3 stomach helminths will increase in spring-summer in southern and central Europe but is expected to proportionately decrease in northern Europe; no major changes are anticipated in the winter months for this group of helminths (Figure S7h).

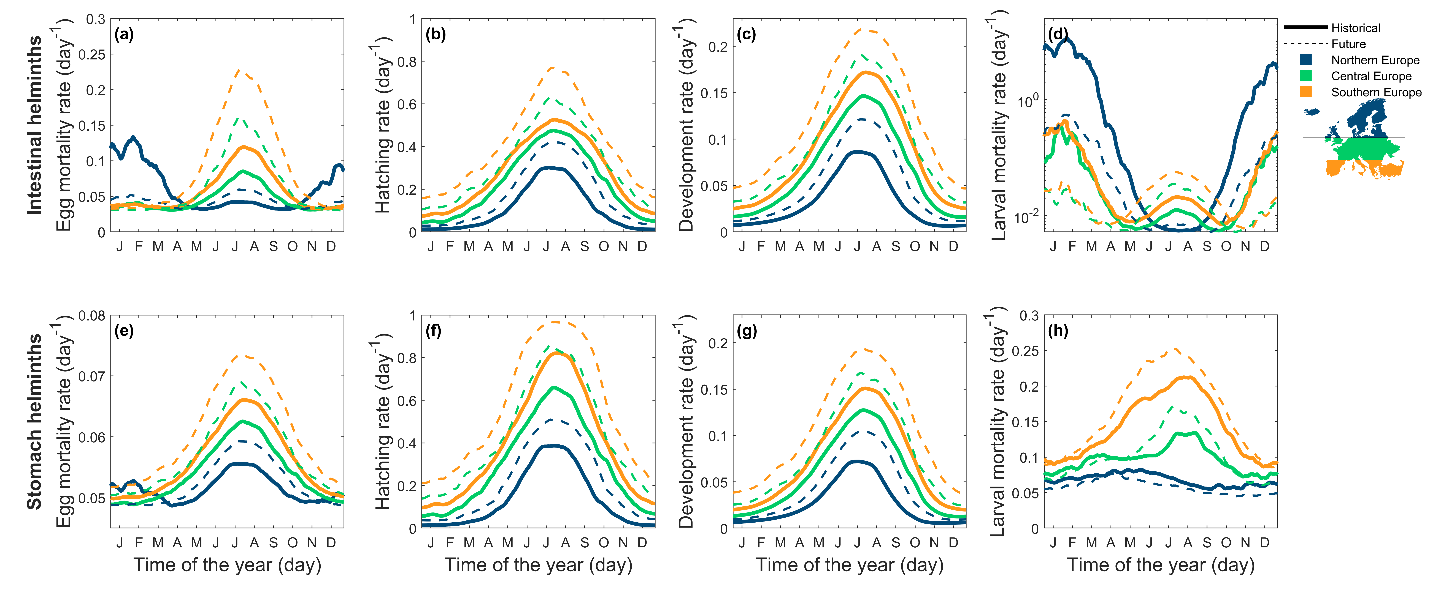

**Figure S7.** Average historical (1981-2000, bold lines) and future (2071-2090 under RCP 8.5 scenario, dashed lines) seasonality of (a,e) egg mortality, (b,f) egg hatching, (c,g) L1 development and (d,h) L3 mortality of (a-d) intestinal and (e-h) stomach helminths in northern (blue), central (green) and southern (orange) Europe, plotted with a two weeks moving average for visual representation.

**SI.7 Spatial patterns of demographic rates**

Here, we report the estimated spatial pattern of the demographic rates in the historical period (1981-2000) and future changes (2071-2090) under the RCP 8.5 ‘business as usual’ climate change scenario (Figure S8, Figure S9).

In the historical period, egg mortality of intestinal helminths was minimum in central Europe at moderate temperatures, while it was quite low for stomach helminths everywhere (Figure S8a,e). Hatching and development rates for both helminth groups follow a north-south trend, with higher rates at lower latitudes (Figure S8b,c,f,g). Larval mortality greatly differs between stomach and intestinal groups with maximum mortality in northern Europe for the first group and in southern Europe for the latter, respectively (Figure S8d,h).

By the end of the century, climate change will cause a drastic increase in egg mortality, up to 60% for intestinal helminths in the Mediterranean area (Figure S9a,e). For both helminth groups, egg hatching and larva development rates will be higher, in particular in Iceland, Scandinavia and the mountainous areas of central and southern Europe (Figure S9b,c,f,g). Larval mortality of stomach and intestinal helminths is expected to undergo similar changes, with a decrease of mortality in northern areas (-100% intestinal helminths and -30% stomach helminths), counterbalanced by an increase in southern Europe (+100% intestinal helminths and +30% stomach helminths) (Figure S9d,h).

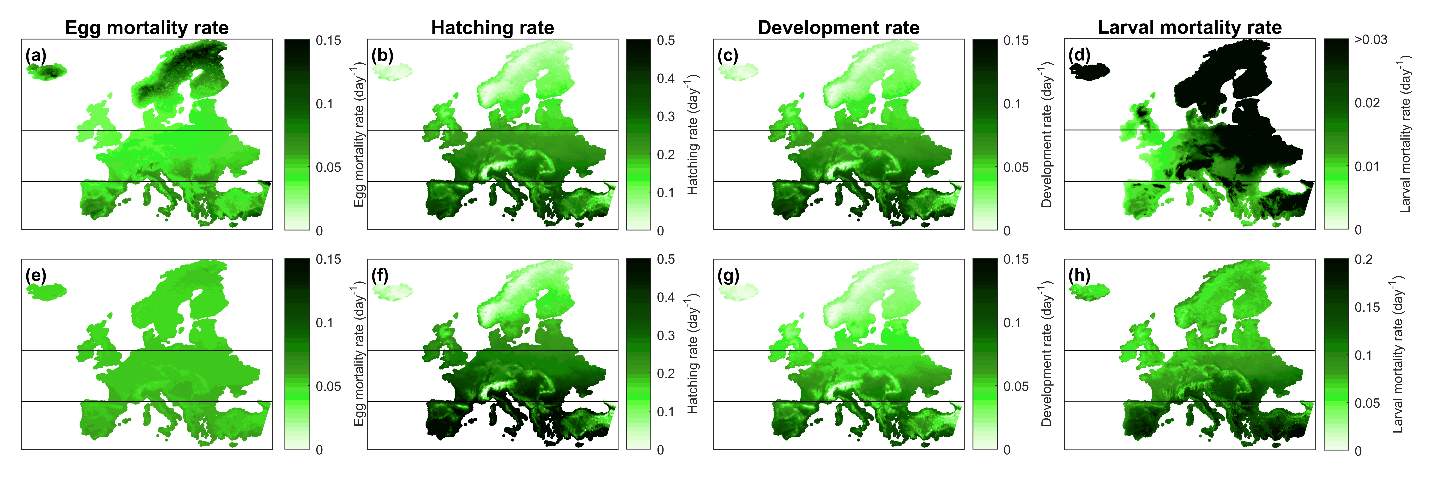

Figure S8. Historical (a,e) egg mortality rate, (b,f) egg hatching rate, (c,g) L1 development rate and (d,h) L3 mortality rate for (a-d) intestinal and (e-h) stomach helminths in the period 1981-2000. Horizontal lines separate northern, central and southern zones.

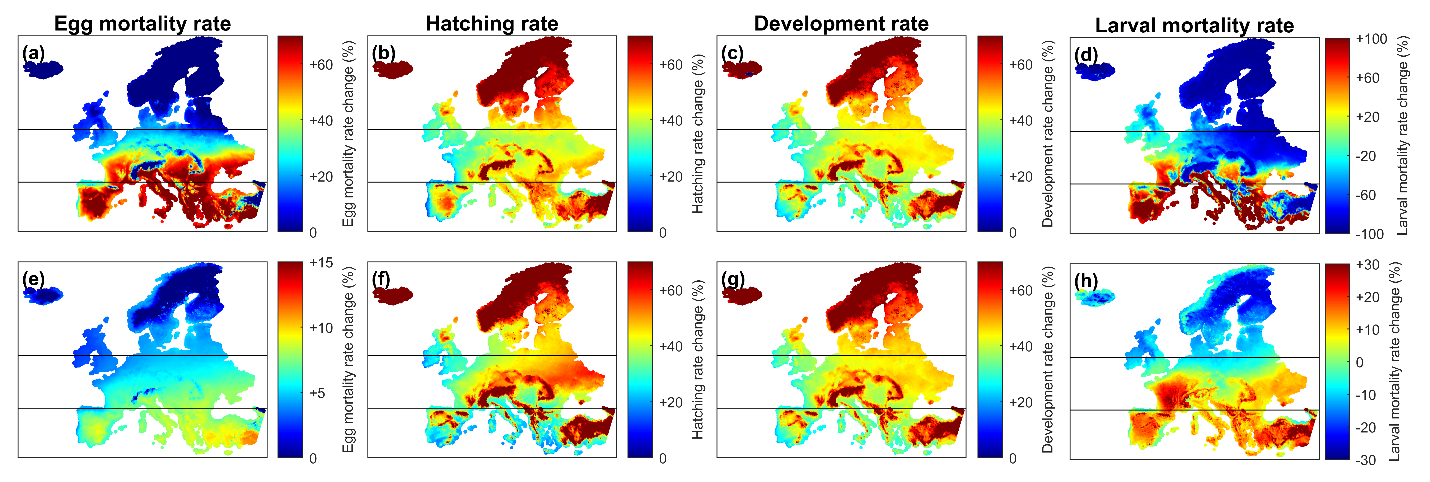

Figure S9. Expected percentage changes of (a,e) egg mortality rate, (b,f) egg hatching rate, (c,g) L1 development rate and (d,h) L3 mortality rate for (a-d) intestinal and (e-h) stomach helminths in the future (2071-2090), compared to the historical period (1981-2000), under the RCP 8.5 scenario. Horizontal lines separate northern, central and southern zones.

**SI.8 Co-occurrence of intestinal and stomach helminths**

We examined the historical distribution (1981-2000) of the infection hazard of intestinal and stomach helminths to classify spatial cells according to their occurrence type (i.e. single occurrence intestinal helminths, single occurrence stomach helminths, co-occurrence, and absence of both groups). We used the 25% percentile threshold of the distribution of each group, $\bar{r}_{intestinal}$=8.69 and $\bar{r}_{stomach}$=6.51 (Figure S10, dashed black lines) to classify each European spatial cell according to the hazard level of the two helminth groups in four infection types: i) single occurrence of intestinal helminths ($r_{intestinal}>\bar{r}_{intestinal}$ and $r_{stomach}<\bar{r}_{stomach}$); ii) single occurrence of stomach helminths ($r_{stomach}>\bar{r}_{stomach}$ and $r_{intestinal}<\bar{r}_{intestinal}$); iii) co-occurrence ($r_{intestinal}>\bar{r}_{intestinal}$and $r_{stomach}>\bar{r}_{stomach}$); iv) absence of both groups ($r_{intestinal}<\bar{r}_{intestinal}$and $r_{stomach}<\bar{r}_{stomach}$). We note that the selected threshold has no direct relationship with the effective abundance of infective stages on the pasture, which could be extrapolated once field data become available.

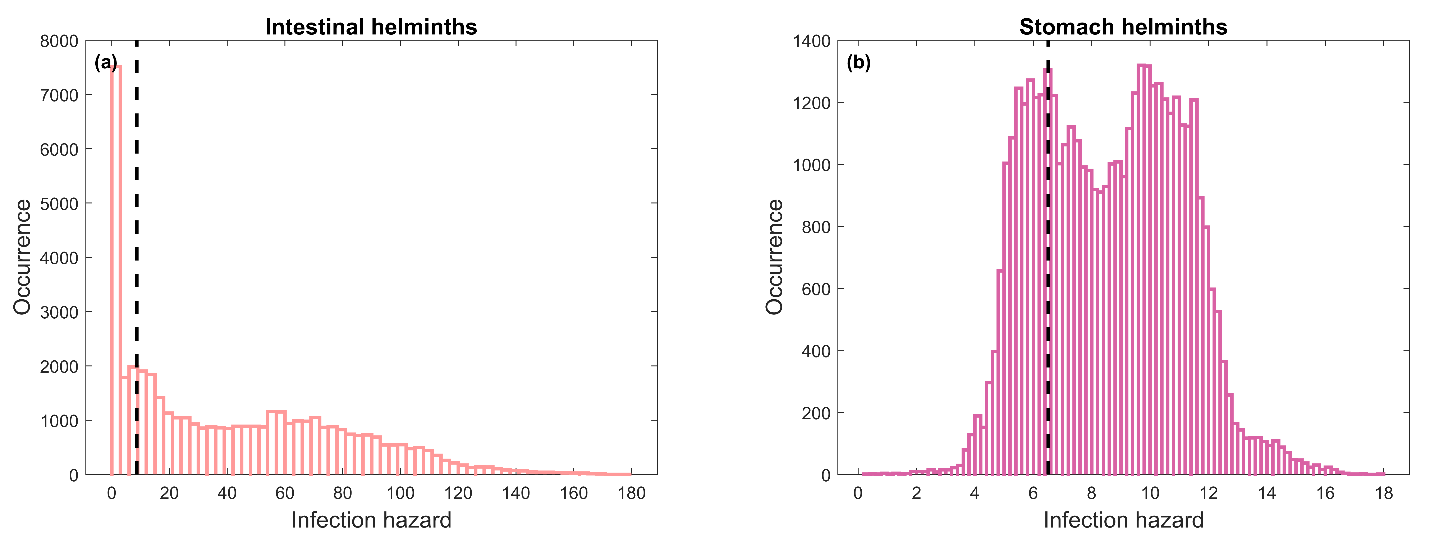

Figure S10. Histogram of the historical (1981-2000) distributions of the infection hazard of (a) intestinal helminths (pink) and (b) stomach helminths (purple) in Europe. Vertical dashed black lines represent the 25% percentile.
